# Ecological analyses of mycobacteria in showerhead biofilms and their relevance to human health

**DOI:** 10.1101/366088

**Authors:** Matthew J. Gebert, Manuel Delgado-Baquerizo, Angela M. Oliverio, Tara M. Webster, Lauren M. Nichols, Jennifer R. Honda, Edward D. Chan, Jennifer Adjemian, Robert R. Dunn, Noah Fierer

## Abstract

Bacteria within the genus *Mycobacterium* can be abundant in showerheads, and the inhalation of aerosolized mycobacteria while showering has been implicated as a mode of transmission in nontuberculous mycobacterial (NTM) lung infections. Despite their importance, the diversity, distributions, and environmental predictors of showerhead-associated mycobacteria remain largely unresolved. To address these knowledge gaps, we worked with citizen scientists to collect showerhead biofilm samples and associated water chemistry data from 656 households located across the U.S. and Europe. Our cultivation-independent analyses revealed that the genus *Mycobacterium* was consistently the most abundant genus of bacteria detected in residential showerheads, yet mycobacterial diversity and abundances were highly variable. Mycobacteria were far more abundant, on average, in showerheads receiving municipal versus well water, and in U.S. households as compared to European households, patterns that are likely driven by differences in the use of chlorine disinfectants. Moreover, we found that water source, water chemistry, and household location also influenced the prevalence of specific mycobacterial lineages detected in showerheads. We identified geographic regions within the U.S. where showerheads have particularly high abundances of potentially pathogenic lineages of mycobacteria and these ‘hot spots’ generally overlapped with those regions where NTM lung disease is most prevalent. Together these results emphasize the public health relevance of mycobacteria in showerhead biofilms. They further demonstrate that mycobacterial distributions in showerhead biofilms are often predictable from household location and water chemistry, knowledge that advances our understanding of NTM transmission dynamics and the development of strategies to reduce exposures to these emerging pathogens.

## Introduction

Bacteria grow and persist in biofilms coating the inside of showerheads and shower hoses despite the seemingly inhospitable conditions found in these habitats. These bacteria must tolerate rapid temperature fluctuations, long intervals of stagnation or desiccation followed by high shear turbulent flow events, and the low nutrient and organic carbon concentrations typical of most drinking water. In many cases, showerhead-associated bacteria must also be able to tolerate residual chemical disinfectants (including chlorinated compounds) which are often added to municipal drinking water to limit bacterial contamination. Despite these stressors, bacterial abundances often exceed 10^6^ cells cm^-2^ inside shower plumbing (1-3). Thus, in the act of showering, we are exposed to elevated concentrations of showerhead-associated bacteria as they are dislodged and aerosolized (4-6).

Most of the bacteria that can become aerosolized and inhaled when the shower is in use are likely harmless. However, this is not always the case. Bacteria within the genus *Mycobacterium* are commonly detected in showerhead biofilms and throughout the water distribution system (7). There are nearly 200 described species of nontuberculous mycobacteria (NTM), which are defined as any members of the genus that are not *M. tuberculosis* or *M. leprae* − which cause tuberculosis and leprosy, respectively. Many species of NTM, including *M. abscessus, M. kansasii*, members of the *M. avium* or *M. fortuitum* complexes, and *M. mucogenicum* (8, 9), are frequently implicated as environmentally-acquired pathogens that can cause an array of human diseases, most notably chronic suppurative lung disease, skin and soft tissue infections often associated with surgical procedures or occupational exposures, cervical lymphadenitis in children, and disseminated infections among susceptible individuals, including those with compromised immune systems (10-12). Although NTM can be found in a range of environments, including soil, house dust, and natural bodies of water, the inhalation of aerosolized NTM from showerheads is frequently implicated as an important route of NTM transmission (4, 13-15). Indeed, previous studies (5, 16-18) have shown that the mycobacterial strains recovered from NTM-infected patients can often be genetically matched to the same mycobacterial strains found in the showers used by the patients.

NTM infections are increasingly recognized as a threat to public health as they are often difficult to treat and the prevalence of NTM lung disease is on the rise in the United States (U.S.) and other developed nations (7, 9, 19). Interestingly, the prevalence of NTM lung disease is not evenly distributed across geographic regions. For example, there are ‘hotspots’ of NTM disease within the U.S. − in Hawaii, southern California, Florida, and the New York City area (20). Moreover, the specific NTM strains most often linked to respiratory infections may vary across geographic regions (19-22). However, it remains unclear why NTM lung disease is increasingly prevalent and why this geographic variation in NTM infections exists. We posit that these patterns are, in part, linked to differences in the amounts and different lineages of mycobacteria found in showerheads.

As mycobacteria are significantly more resistant than other bacteria to chlorine and chlorine byproducts (18, 23, 24), they are expected to be more abundant in showerheads and water distribution systems where such disinfectants are used. We also expect some mycobacteria to be more common in households receiving more acidic water (18, 25, 26), and in water systems where free-living amoebae (FLA) are common, given that mycobacteria can survive and replicate inside amoebae (27-30). However, it remains unclear whether these abiotic and biotic variables can effectively predict the distributions of NTM in household water distribution systems and the likelihood of acquiring NTM opportunistic infections.

The diversity, distributions, and ecologies of those NTM colonizing showerheads are clearly relevant to public health. Unfortunately, we currently lack a comprehensive understanding of which mycobacteria are found in showerheads and how the distributions of mycobacterial taxa vary depending on geographic location, interactions with other microorganisms, and environmental conditions. These are the knowledge gaps that motivated this study − we sought to understand the geographic and environmental factors that predict the amounts and types of NTM found in showerhead biofilms. Likewise, given the potential for these bacteria to cause respiratory disease, we investigated whether there is a concordance between the geographic distributions of showerhead mycobacteria with the prevalence of NTM lung disease, the latter estimated from two separate epidemiological surveys.

## Results and Discussion

### Mycobacteria abundance across showerheads

We worked with citizen scientists to collect showerhead biofilm samples from locations across the U.S. and Europe, with 656 samples included in downstream analyses (606 in the U.S., 50 in Europe, see Figure S1). We extracted DNA directly from these biofilm samples and used a cultivation-independent 16S rRNA gene sequencing approach to assess overall bacterial community composition in each sample. We found that bacteria assigned to the genus *Mycobacterium* were, on average, the most abundant group of bacteria found in the showerhead biofilm samples (Figure S2). Other abundant taxa included members of the genera *Sphingomonas, Bradyrhizobium, Blastomonas*, and *Phenylobacterium*, bacterial taxa commonly observed in water distribution systems (1, 15, 31). Across all samples, the mean abundance of the *Mycobacterium* genus was 13.5% of 16S rRNA gene reads (Figure S2). However, the abundance of mycobacteria was highly variable across samples, ranging from 0% (no mycobacteria detected) to >99% of bacterial 16S rRNA gene reads, with *Mycobacterium* accounting for >10% of 16S rRNA gene reads in 238 of the total 652 samples. The observed ubiquity of mycobacteria is comparable to other studies that have used cultivation-independent methods to assess mycobacterial abundances in water distribution systems (4, 14, 15, 31).

Overall, mycobacteria were more than twice as abundant in U.S. homes receiving water from municipal water treatment plants as compared to homes on well water (P < 0.001; Figure 1A). Water treatment plants in the U.S. must maintain excess chlorine-based disinfectant within the distribution system (32) and, as expected, we found that those homes on municipal water had measured total chlorine concentrations 15 times higher, on average, than homes with well water (Figure S3). The observed higher abundances of mycobacteria in showerheads receiving municipal versus well water is consistent with previous work (15) and is likely a result of disinfection selecting for mycobacteria as they are typically more resistant than other bacteria to the toxic effects of chlorine and chloramine (24, 31, 33). We also found that the relative abundances of mycobacteria in plastic showerheads were, on average, two times lower than in showerheads that were either metal or a mix of metal and plastic components (P<0.01, Figure 1B). Similar results have been reported previously (34, 35) and these patterns are likely a product of the leaching of biodegradable carbon from plastic materials supporting elevated growth of other bacterial taxa that can outcompete mycobacteria in showerhead biofilms (1, 36).

**Figure 1:**
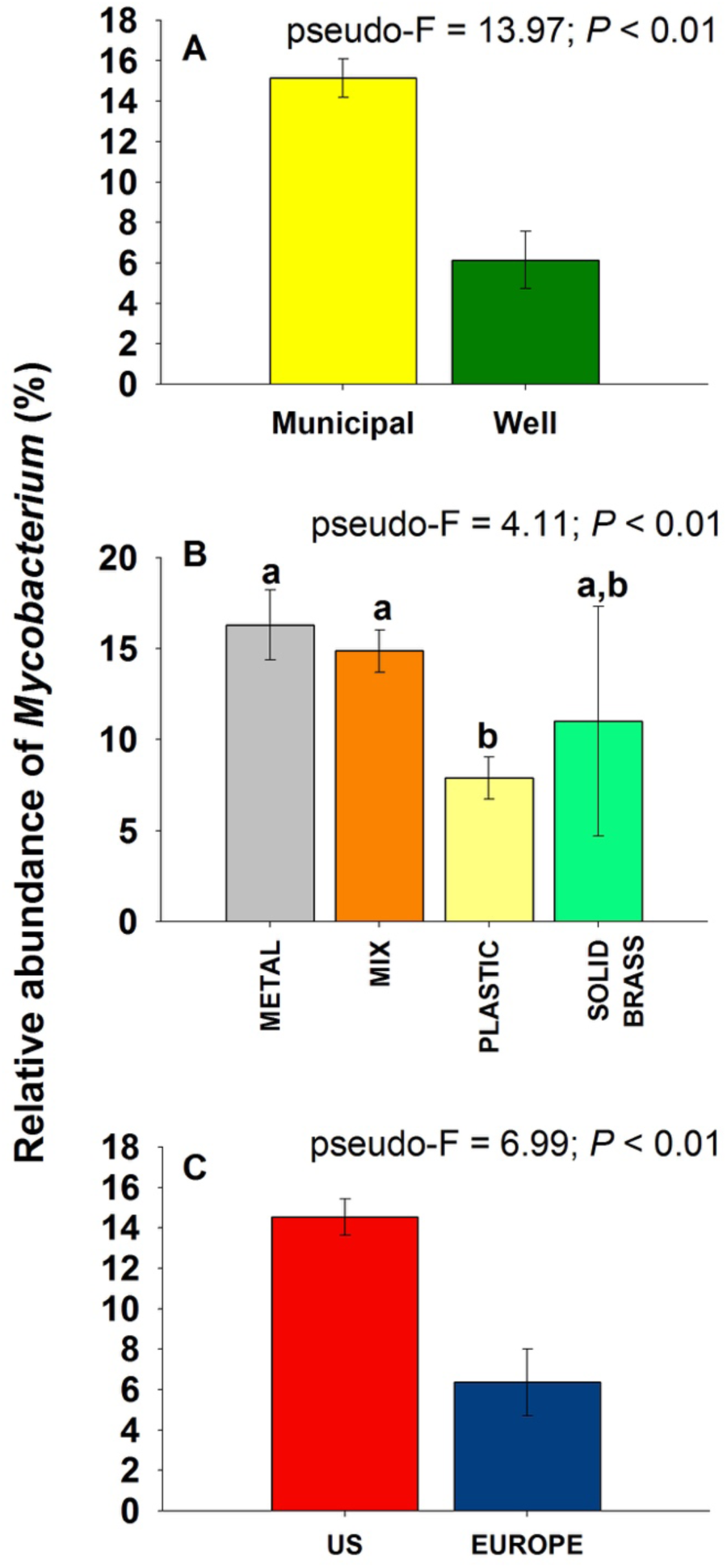
Differences in the relative abundance of mycobacteria (as determined via 16S rRNA gene sequencing) across households in the U.S. on municipal versus well water (A), across showerheads constructed of different materials (B), and across households in the U.S. versus Europe (C).

It has been suggested that mycobacteria should be less abundant in biofilm samples dominated by methylotrophs, particularly *Methylobacterium* spp. (37, 38). In contrast to this expectation, we found that showerheads with higher abundances of *Methylobacterium* did not necessarily have lower proportional abundances of mycobacteria (Figure S4). Therefore, *Methylobacterium* is unlikely to be a useful indicator of mycobacterial loads. We did, however, identify a positive correlation between mycobacterial abundances and the abundances of free-living amoebae (FLA) within the Vermamoebidae group in a subset of samples (N = 89) for which we obtained both bacterial and eukaryotic small-subunit rRNA gene sequence data (Pearson's r = 0.25, *P* = 0.02). This pattern was mainly driven by *Vermamoeba* (*Hartmannella*) *vermiformis*, a species of amoebae that was the most abundant microeukaryote detected in our samples and commonly found in water distribution systems (30, 39) with this species previously shown to be capable of harboring intracellular mycobacteria (28, 29). The apparent co-occurrence of FLA and mycobacteria highlights the importance of considering potential associations between protists and bacteria when trying to predict distributions of mycobacteria and other potentially pathogenic bacteria (e.g. *Legionella*) in residential water systems (40-42).

The genus *Mycobacterium* was not equally abundant across all geographic regions included in our sampling effort (Figure S5). Strikingly, showerhead biofilms in the U.S. had, on average, 2.3 times higher abundance of taxa assigned to the genus *Mycobacterium* as compared to showerheads in Europe (P <0.01, Figure 1C). The lower abundances of mycobacteria in Europe as compared to the U.S. may be driven by differences in water treatment and water distribution systems (23). Indeed, analyses of shower water chemistry by the citizen scientists support this hypothesis as we found that U.S. households receiving municipal water had significantly higher chlorine and iron concentrations, but significantly lower pH and nitrate levels, than European households on municipal water (Figure S6). Most notably, total chlorine concentrations in the U.S. were 11 times higher, on average, than shower water measured in the European households (Figure S6). The practice of adding chlorine-based residual disinfectants during water treatment is less common in Europe than in the U.S. (43) and this key difference in water treatment practices may be contributing to the elevated abundances of mycobacteria observed in showerheads from U.S. households.

### Mycobacterial diversity in showerheads

The aforementioned analyses focused on measured abundances of the genus *Mycobacterium*, as inferred from 16S rRNA gene sequence data. However, the genus *Mycobacterium* includes nearly 200 species that can differ with respect to their ecologies and pathogenicity (44). Thus, to obtain more detailed information on what specific mycobacteria, including potential pathogens, were found in the showerhead biofilms, we amplified DNA from each of the 656 showerhead samples and sequenced a portion of the hsp65 (65 kDa heat shock protein) gene using mycobacteria-specific primers (see Methods). In total, we recovered 1,029 mycobacterial exact sequence variants (ESVs) with the 100 ESVs shown in Figure 2 accounting for >95% of the reads. These ESVs span nearly the full extent of known mycobacterial diversity, highlighting that biofilms of household water distribution systems can harbor an extensive array of mycobacterial species, including a number of taxa that are not closely related to any previously described isolates (Figure 2). We combined the top 100 ESVs into 34 phylogenetically-defined clades for downstream analyses. We found that the dominant mycobacterial clades recovered from the showerhead biofilms included potential pathogens that are frequently recovered from patients diagnosed with NTM infections (e.g. *M. avium* complex, *M. abscessus, M. fortuitum* complex) as well as mycobacteria that are not typically considered pathogenic (e.g. *M. gordonae, M. hassiacum, M. canariasense*) (Figure 2). Information on the ubiquity and median abundance of the dominant mycobacterial clades detected in the showerheads using our cultivation-independent hsp65 sequencing approach are provided in Figure 2 and Figure 3, respectively.

**Figure 2:**
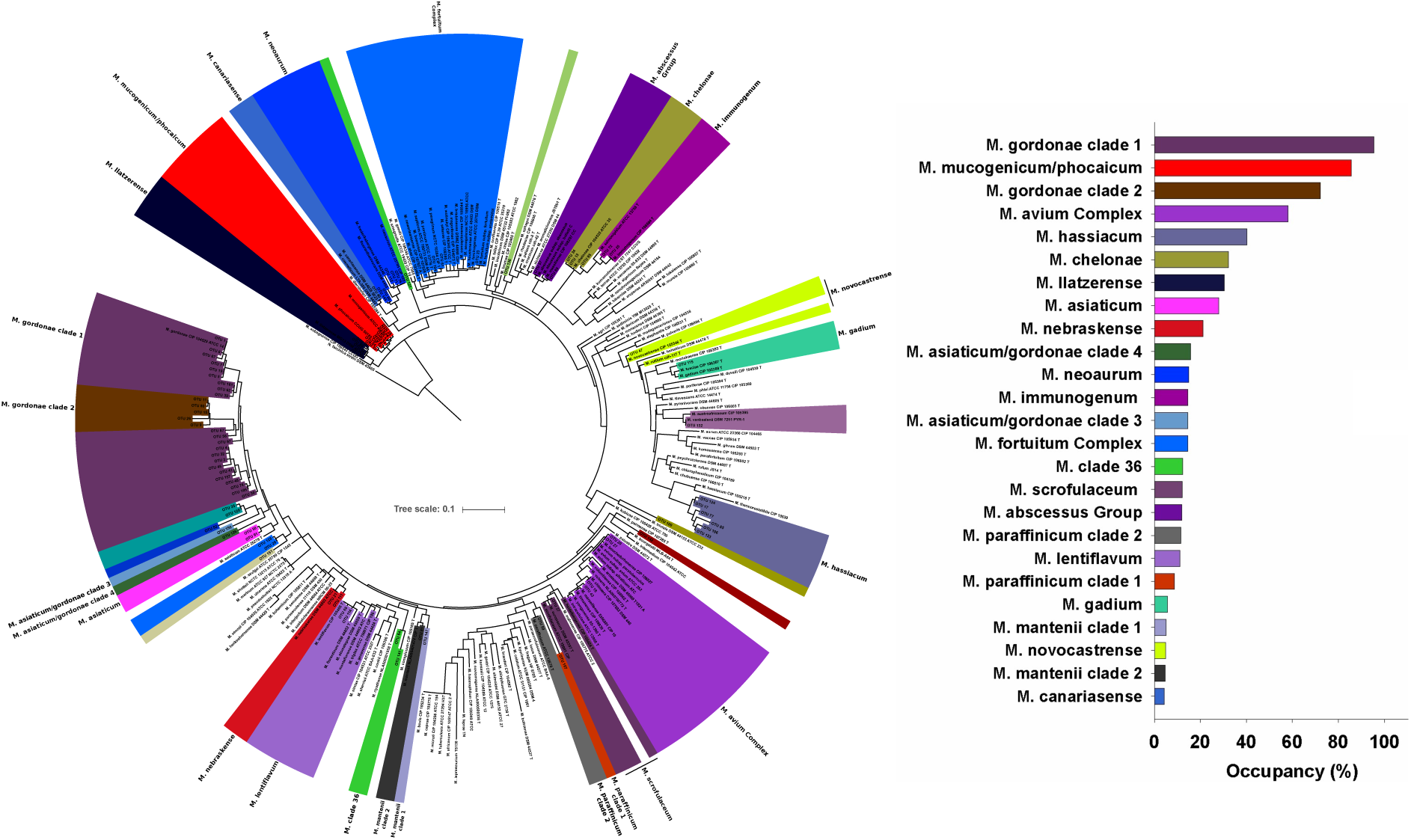
Phylogenetic tree showing the mycobacterial diversity recovered from the cultivation-independent analyses (hsp65 gene sequencing). Included in the tree are those reference mycobacterial strains from Dai et al. (59). The colors indicate the 34 cleades of *Mycobacteria* with the labels indicating the taxonomic identity of each clade. The tree was rooted with a hsp65 sequence from *Nocardia farcinica* (DSM43665). The plot on the right shows the percent occupancy of the top 25 mycobacterial clades with occupancy assessed as the percentage of samples (out of 656 in total) in which each clade was detected. Colors indicate unique mycobacterial clades with the color scheme used in the tree matching the color scheme used in the associated plot of mycobacterial occupancy.

**Figure 3:**
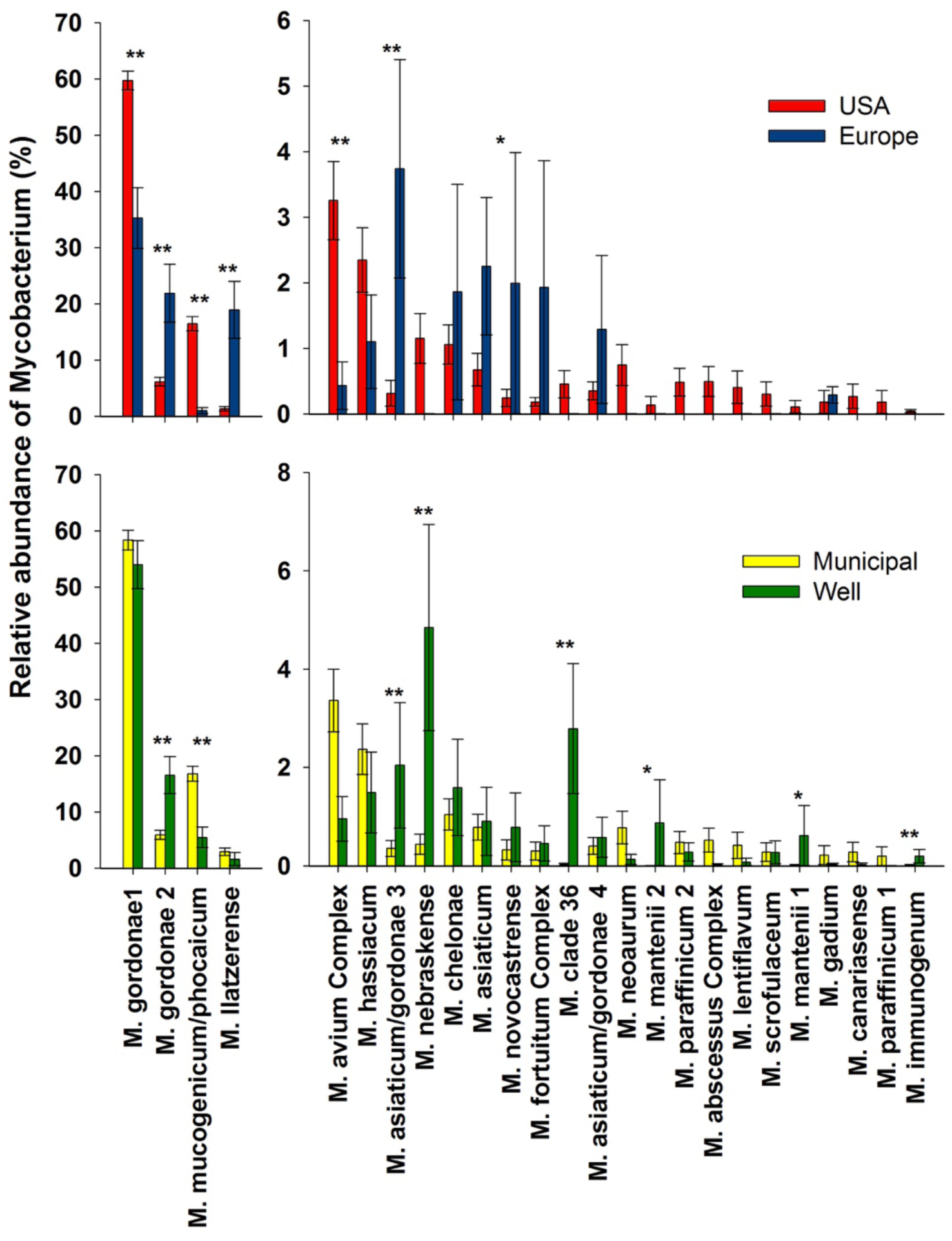
Abundances of the top 25 mycobacterial clades detected across homes in the U.S. on well versus municipal water (N = 520 and 86, respectively, panel A) and across homes in the U.S. versus Europe (municipal water only, N = 606 and 50, respectively, panel B). The y-axes were split to better illustrate the differences among the less-abundant clades.

Most of the mycobacterial diversity found in the showerhead biofilm samples was not captured in a corresponding cultivation-dependent survey. For those 186 samples which were analyzed using both cultivation-dependent and cultivation-independent approaches, we identified nearly four times more mycobacterial ESVs with the cultivation-independent approach (Figure S7). Many clades (including some of the more abundant clades, such as *M. gordonae, M. hassiacum*, and *M. llatzerense*) were missed completely with the cultivation-dependent approach (Figure S7). While some mycobacterial clades were detected with both approaches (Figure S7), culturing captured only a small fraction of the total mycobacterial diversity found in showerheads. These findings confirm results from previous studies (26, 45) that many mycobacteria in the environment (and possibly in patients with respiratory infections) are simply missed when using standard culture techniques. Our results highlight the importance of using cultivation-independent approaches for detecting mycobacteria when possible, as mycobacterial diversity in these biofilm samples, and other sample types, is likely to be significantly underestimated when relying on cultivation-based surveys.

### Biogeography of selected mycobacteria

The individual mycobacterial clades detected by our cultivation-independent hsp65 gene sequencing analyses, including clades with known pathogens, often exhibited distinct geographic patterns. Not all mycobacterial lineages were likely to be found everywhere − the mycobacterial diversity found in showerheads varied as a function of household location (Figures 3-4; and S8-S9). Some clades were far more abundant in Europe than in the U.S., including multiple clades related to *M. gordonae* and a clade that included *M. llatzerense*, which was 14 times more abundant in Europe than in the U.S. (Figure 3). We note that a study of mycobacteria in Parisian tap water systems also found *M. llatzerense* to be dominant, even though this species is rarely detected in the U.S. (23, 46). In contrast, we found mycobacteria within the *M. avium* complex (MAC), that includes multiple opportunistic pathogens, were relatively more abundant in U.S. showerheads than in those from Europe. Although these differences in the abundances of specific mycobacterial groups between U.S. and European households may be a product of dispersal limitation, we expect that these patterns are more likely driven by differences in water chemistry (Figure S6), water distribution systems, or water treatment practices. Although showerheads are only one potential source of NTM infections, the significant differences in the mycobacterial communities inhabiting U.S. versus European showerheads may explain some of the documented geographic variation in the clinical isolates obtained from NTM patients on the two continents (22).

**Figure 4:**
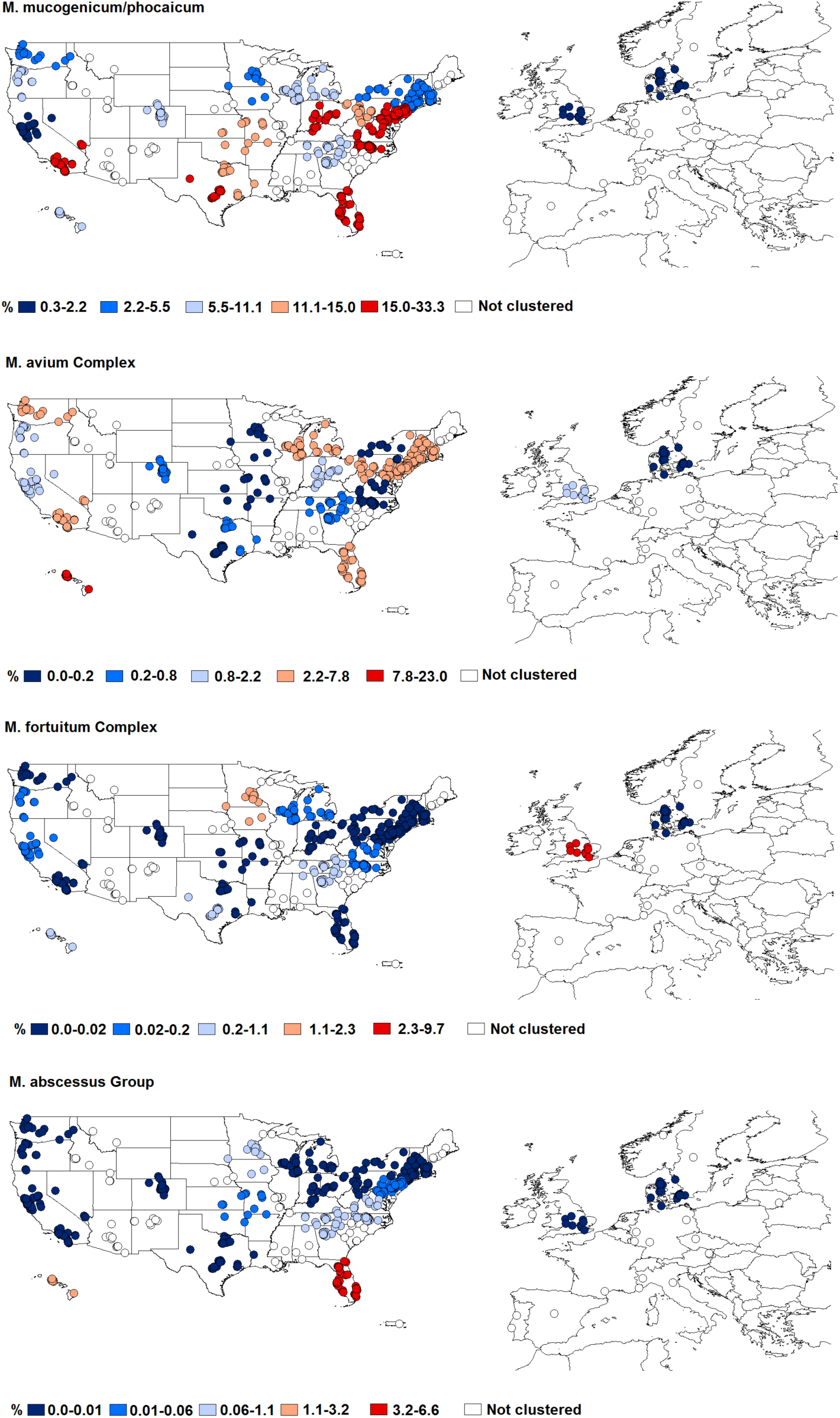
Differential abundances of each of four lineages of mycobacteria that include pathogens across the geographic clusters of showerhead biofilm samples. The different colors indicate the mean abundances of each mycobacterial lineage across each of the 23 geographic clusters identified (uncolored points are samples from households not included in any of these 23 clusters). For details on each cluster and the abundances of mycobacteria within each cluster, see Figure S8.

Within the U.S., households supplied by municipal versus well water had distinct mycobacterial communities (Figure 3). Most notably, homes on municipal water had higher abundances of the *M. mucogenicum/phocaicum* clade, potentially pathogenic mycobacteria (47). Other mycobacterial taxa, including *M. nebraskense* and some *M. gordonae* clades, were more abundant in U.S. homes that use well water. However, these taxa are rarely considered pathogenic (10, 12, 48) suggesting that exposures to pathogenic mycobacteria from showerheads are typically higher in homes on municipal water. This finding is consistent with reports that water source has been found to play a role in NTM disease prevalence (24, 49).

Some of the mycobacterial lineages that include known pathogens were more likely to be abundant in showerheads from certain regions of the U.S. (Figure 4, Figures S8-S9). We found that four mycobacterial clades that are often considered pathogenic (*M. mucogenicum/phocaicum*, the *M. avium* complex, the *M. fortuitum* complex, and the *M. abscessus* group) exhibited significant geographic variation in their abundances across the U.S. (Figure 4). While these four pathogenic clades exhibited distinct geographic patterns, showerheads from homes in Hawaii, southern California, Florida, the upper Midwest, and the mid-Atlantic states consistently had higher abundances of one or more of these pathogenic mycobacteria. Most notably, the *M. abscessus* group and *M. avium* complex were far more abundant in Hawaii, Florida, and the northeastern U.S. than in other regions. The geographic ‘hot spots’ in mycobacterial abundances shown in Figure 4 were, to some degree, predictable from shower water chemistry. The abundances of these pathogenic mycobacterial lineages in showerheads were often significantly correlated with mean annual outdoor temperature and shower water total chlorine, alkalinity, and pH levels, with the specific direction of these correlations depending on the clade in question (Figure S10). Most notably, showerheads found in households from warmer locations with higher shower water chlorine concentrations were found to have higher abundances of the *M. mucogenicum/phocaicum* clade. However, much of the observed geographic variation in the abundances of other mycobacterial groups (including the *M. abscessus* group and the *M. avium* complex) could not be reconciled from the variables included in our model (Figure S10). This may be because we did not measure all of the relevant shower water characteristics (e.g. organic carbon concentrations, water heater temperature) or because the conditions that favor the growth of these mycobacteria are determined by source water characteristics or conditions at other points in the water distribution network. Nevertheless, we did find that individual mycobacterial lineages have distinct geographic distributions and environmental preferences, information that can be used to improve predictions of which showerheads are most likely to harbor pathogenic mycobacteria.

### Showerhead-associated mycobacteria and the epidemiology of NTM lung disease

Those regions within the U.S. that we identified as having relatively high abundances of known mycobacterial pathogens in showerhead biofilms (Figure 4) generally overlapped with those regions in the U.S. previously reported as having a higher-than-average prevalence of NTM lung disease (20). For example, some of the U.S. regions with the highest reported incidences of patients diagnosed with NTM lung disease (including Florida, Hawaii, southern California, and the mid-Atlantic states) are the same regions from which we found showerheads with high abundances of potentially pathogenic mycobacteria (Figure 4). To further investigate this pattern, we explicitly tested whether the distributions of pathogenic NTM across the U.S. showerheads in this study were correlated with geographic patterns in NTM lung disease prevalence. We compared the state-level median abundances of two potentially pathogenic mycobacterial clades that were frequently detected in showerheads (*M. abscessus* group and *M. avium* complex) against the reported prevalence of NTM lung disease across Medicare beneficiaries and persons with cystic fibrosis (Figure S11). We expected only a modest correlation, if any, given the relatively small number of showerheads per state and the constraints associated with accurately estimating NTM lung disease prevalence (20, 50). However, we found significant geographic correlation between the abundances of these mycobacteria found in showerheads sampled across the U.S. and NTM lung disease prevalence, the latter derived from two independent datasets (Pearson’s r >0.6, P<0.001 in both cases, Figure S11). These results add to the growing body of evidence that mycobacteria living in showerheads are likely an important source of NTM infections.

## Conclusions

Mycobacteria are frequently abundant in showerheads and many showerheads harbor mycobacterial lineages that include known pathogens. The amount and types of mycobacteria found in showerheads appear to vary, depending on household location, water source, water chemistry, the presence of free-living amoebae, and showerhead material. Moreover, those geographic regions with higher abundances of mycobacterial pathogens tend to be ‘hot spots’ of NTM lung disease within the U.S. So far, our results are correlative, but they are in line with previously-published work on the factors structuring mycobacterial abundances and the importance of showerheads as a source of NTM infections. We also demonstrate the synergy of coupling an extensive citizen scientist sampling effort with molecular diagnostic techniques to comprehensively investigate the factors associated with the likelihood of acquiring NTM lung disease. More generally, our results highlight the relevance of understanding how shifts in household water sources (i.e., from well to municipal water sources) and water treatment practices may be contributing to the apparent rise in NTM infections in U.S. and European populations.

## Materials and Methods

### Sample collection

Showerhead biofilm samples were collected by citizen scientists participating in the Showerhead Microbiome Project (goo.gl/7G6xbc). We recruited participants using the website, social media, and email campaigns from throughout the U.S. and Europe from July 2016 to November 2016. Enrolled participants were provided a written Informed Consent form approved by the North Carolina State University’s Human Research Committee (Approval No. 9158). Each participant was then provided with a sampling kit that contained a dual-tipped sterile Puritan CultureSwab, a water chemistry analysis kit, sterile gloves to be worn during sampling, and a brief questionnaire. Participants were not queried about their NTM infection status. All biofilm samples were collected by swabbing the interior of an unscrewed showerhead, with participants asked to swab the most commonly-used showerhead in each household, as close to the inside interface of the showerhead, as possible. Each participant was asked to provide the household address, the household water source, the estimated time since installation of showerhead, its usage frequency, cleaning frequency, and a description of the showerhead sampled (including materials and spray pattern). All swab samples collected from the U.S. were mailed directly to the University of Colorado where they were stored in a −20°C freezer until processed. Swab samples from Europe were mailed directly to Copenhagen, Denmark where they were stored at −20°C until the European collection was completed. The European swab samples were then sent overnight on dry ice to the University of Colorado where they were stored at −20°C until processed. In total, we collected 691 samples, 638 showerhead biofilm samples from across the U.S. (49 of 50 U.S. states) and 53 samples from Europe (13 different countries) − see Figure S1.

### Water chemistry analyses

Each participant conducted basic chemical analyses of the water collected from the same shower used for the biofilm sample collection. Water chemistry was determined for each showerhead using Hach Aquachek Water Quality Test Strips (Hach, Loveland, CO, USA). Each kit included a *‘5-in-1’* test strip (which measures total chlorine, free chlorine, total hardness, total alkalinity, and pH), a test strip for nitrate and nitrite, and a test strip for total iron concentrations.

### 16S rRNA gene sequencing to characterize showerhead bacterial communities

We used an approach described previously (51) to amplify and sequence the V4 hypervariable region of the 16S rRNA gene from all 691 biofilm samples. DNA was extracted from one of the two swabs collected per showerhead using the Qiagen PowerSoil DNA extraction kit and then PCR amplified in duplicate reactions using the 515f/806r primer pair modified to include Illumina adapters and the appropriate error-correcting barcodes (52). Each 25 μL reaction included 12.5 μL of Promega HotStart Mastermix, 10.5 μL of PCR-grade water, 1 μL of PCR primers (combined at 10 μM), and 1 μL of purified genomic DNA. The thermocycler program was an initial step at 94°C for 3 min, following by 35 cycles of 94°C for 45 s, 50°C for 1 min, and 72°C for 1.5 min. The program concluded with a final elongation step at 72°C for 10 min. Both ‘no template’ controls and ‘DNA extraction kit’ controls were included with each set of 90 samples to check for potential contamination. Duplicate reactions were pooled, cleaned, and normalized using the ThermoFisher SequalPrep Normalization Plate kit. Amplicons were sequenced on 3 MiSeq runs at the University of Colorado Next-Generation Sequencing Facility with the 2×150 bp paired-end chemistry.

After demultiplexing and combining the data from all 3 sequencing runs, paired reads were merged with a minimum length of 200 bp for the merged sequence, and quality filtered using the uSearch10 pipeline (53), discarding sequences with greater than 1 error per base call. The quality-filtered reads were processed using uNoise3 (54) to identify exact 16S rRNA gene sequence variants (ESVs). Taxonomy was determined for each ESV using the Ribosomal Database Project Classifier (55) trained on the Greengenes database (56). After removing those ESVs that were represented by <25 reads across the entire dataset and those classified as mitochondria or chloroplasts, we ignored any samples that yielded less than 2,000 reads per sample. 39 of the 691 biofilm samples and all of the ‘no template’ and ‘DNA extraction kit’ negative controls failed to meet this threshold for sequencing depth and were excluded from downstream analyses. The percentage of mycobacteria in each sample was calculated by summing up all reads classified to the *Mycobacterium* genus.

### Mycobacterial-specific hsp65 sequencing

As the 16S rRNA gene sequencing analyses described above provide insufficient resolution to differentiate between many mycobacterial lineages, we also analyzed all biofilm samples by PCR amplifying and sequencing a region of the hsp65 gene using mycobacterial-specific PCR primers (57), including negative controls in all analyses as described above.

A ‘two-step PCR’ protocol was used for the hsp65 amplifications with sample-specific barcodes ligated to the hsp65 amplicons in a second round of PCR. The first PCR step was conducted with duplicate reactions per extracted DNA sample using the Tb11-Tb12 primers (57) that included the appropriate Illumina adapters. Each 25 μL PCR recipe was identical to that described above with the following thermocycler program: a 3 min initial step at 94°C, followed by 45 cycles of 94°C for 60 s, 60°C for 60 s, and 72°C for 60 s, with a final 10 min elongation step. Duplicate reactions were pooled and the amplicons were cleaned using the Qiagen UltraClean PCR Clean-Up Kit following the manufacturers’ instruction.

A second round of PCR was then performed to attach a unique 12-bp error-correcting barcode to the amplicons from each sample to allow for multiplexing. Each of the 42 μL reactions included 20 μL of Promega Hotstart Mastermix, 14 μL of PCR-grade H_2_O, with 4 μL of the forward/reverse universal 12-bp barcodes (10 μM each) and 2 μL of the Tb11-Tb12 amplicons from the previous round of PCR reactions added to 36 uL of the Mastermix and H_2_O mixture. The thermocycler program included an initial step at 95°C for 3 min, followed by 8 cycles of 95°C for 30 s, 55°C for 30 s, and 72°C for 30 s, with a final step at 72°C for 5 min. The resulting barcoded hsp65 amplicons were then cleaned and normalized using the method described above. Amplicons were sequenced on 2 Illumina MiSeq runs using the 2×300 bp paired-end chemistry at the University of Colorado Next-Generation Sequencing Facility.

After demultiplexing and trimming the reverse reads to 250 bp using fastq_filter, the paired reads were merged with a minimum length of 200 bp for the merged sequence and quality filtered using the uSEARCH10 pipeline (53), discarding those sequences with >1 error per base call. We used the uNOISE3 pipeline (58) to identify exact sequence variants (hsp65 ESVs) with the taxonomy assigned using the Ribosomal Database Project Classifier (55) trained on the hsp65 reference database described in Dai et al. (59). The representative sequences for each ESV were then compared to those 157 hsp65 sequences in the reference database compiled by Dai et al. (59) using the BLAST algorithm (60). Those ESVs that did not have >90% sequence similarity to those in the reference database were removed as they were not from members of the *Mycobacterium* genus. Across the whole dataset, this process removed 5% of the quality-filtered hsp65 reads with most of the removed reads classified as belonging to other genera within the Actinomycetales order. Only those samples with >200 quality-filtered mycobacterial hsp65 reads per sample were included in downstream analyses and this threshold removed 5 of the 691 biofilm samples and all of the ‘no template’ and ‘DNA extraction kit’ negative controls.

We focused our downstream analyses on the top 100 mycobacterial ESVs across the whole mycobacterium filtered dataset (these ESVs accounted for 96% of the quality-filtered mycobacterial hsp65 reads). The 100 representative hsp65 ESV sequences were aligned against the hsp65 reference available in the Dai et al. (59) database using MUSCLE v.3.8.31 (61) and a phylogenetic tree was constructed using RAxML (62) with a hsp65 sequence from *Nocardia farcinica* (DSM43665) to root the tree. All of the ESVs in the best scoring phylogenetic tree were clustered into discrete clades using RAMI (63) with the patristic distance threshold set to 0.05 (clades defined as having >5% patristic distance across the sequenced portion of the hsp65 gene). The taxonomic identity of each clade was determined based on sequence similarity to the type strains that fell within each clade and the phylogenetic tree as visualized using iTOL (64). Information on each of the 100 hsp65 ESVs, their assigned clades, phylogenetic placement and their similarity to type strains are included in Figure 3.

### Protistan analyses

As mycobacteria are known to associate with various protists, we wanted to test for co-occurrence patterns between mycobacteria and specific protists. To do so, we selected a subset of 186 samples for which we obtained 18S rRNA marker gene data following protocols as described in Ramirez el al., 2014 (65). In brief, we used the 1391f/EukBr primer set and pooled and sequenced amplicons along with appropriate controls, as described above for 16S rRNA communities. Reads were demultiplexed, merged and trimmed to 100 bp reads, and quality filtered as described above. A database of ≥97% similar sequence clusters was constructed using USEARCH (53) and taxonomic assignments were made with the PR2 database (66). Of the 186 samples, we retained 89 samples for downstream analyses with a minimum threshold of 1,200 reads per sample, after removing non-microbial eukaryotes including Streptophyta and Metazoa. To account for differential read coverage, we rarefied samples to 1,200 reads per sample for the 18S rRNA analyses and also rarefied the 16S samples to 1,500 reads per sample for the subset of 89 samples for which we obtained protist data.

For the subset of samples that we obtained both 16S and 18S rRNA gene data for (n=89), we investigated whether the relative abundance of the most dominant groups of free living amoebae (FLA) were correlated with mycobacterial relative abundances (at the genus level). To do this, we conducted Pearson’s correlations to evaluate the correlation between the three most abundant FLA families (Vermamoebidae, Acanthamoebidae, and Echinamoebidae) with the relative abundance of the genus *Mycobacterium* across the 89 samples.

### Cultivation-based analyses of mycobacteria

In addition to the cultivation-independent DNA sequencing-based analyses described above, 186 randomly-selected biofilm samples were cultured using a standard approach for the culture and isolation of environmental mycobacteria (36). Briefly, swabs were immersed in 2 mL of autoclaved ultrapure water and vortexed on high for 1 min. To select for mycobacteria, 450 μL of sample was transferred to a sterile Eppendorf tube with 50 μL of 1% cetylpyridinium chloride, vortexed, and incubated at room temperature for 30 min. After incubation, 100 μL of each sample was plated on duplicate Middlebrook 7H10 agar with oleic acid/glycerol enrichment and incubated at 37°C for 21 days. Out of the 186 samples cultured, 40 were positive for mycobacteria (no mycobacterial isolates were recovered from the remaining 146 samples). Thus, we were able to obtain mycobacterial isolates from 40 samples, with these 40 samples yielding 74 unique isolates. We extracted DNA from all 74 isolates, in addition to a ‘blank’ control sample of the media used to grow the isolates, using the Qiagen Powersoil DNA Extraction kit, and sequenced the amplified hsp65 to identify each isolate using the approach described above. All isolates had >98% sequence similarity matches over the entire length of the amplicon to those sequences in the Dai et al. (59) database. The hsp65 sequences from these isolates were placed into the phylogenetic tree described above that included the reference sequences and the sequences obtained from the cultivation-independent analyses of all the biofilm samples. We then assigned the isolate sequences to the respective lineages and determined their taxonomic identity using the methods described above. Information on each of these isolates, their taxonomic identities, and their phylogenetic placement is provided in Figure S7.

### ‘Storage’ study

We conducted a separate experiment to determine how prolonged storage at room temperature affects the showerhead biofilm bacterial communities. We did this to determine if shipping samples unrefrigerated from across the U.S. to Boulder, Colorado may have influenced our determination of mycobacterial relative abundances. For this experiment, we collected 12 replicate swabs placed in individual showerheads at each of 7 households (2 in Colorado, 2 in Hawaii, 2 in North Carolina, 1 in southern California, USA). These swabs were cut 2.5 cm from the tip and placed into the accessible faceplate of 7 ‘polished brass’ showerheads provided by Shower Clear (West Orange, NJ, USA). Participants were asked to replace the pre-existing showerheads in each of the 7 households for 30-40 days. After this time period, the replicate swabs from each showerhead were shipped to the University of Colorado overnight at 4°C, where they were either frozen at −20°C immediately (day 0) or held at room temperature conditions inside sealed bags to minimize drying for 3, 7, and 10 days prior to freezing at −20°C. Genomic DNA was extracted from all swabs (7 homes x 12 replicate swabs per home, 3 per storage duration) and the V4 region of the 16S rRNA gene was amplified and sequenced from the extracted DNA using the protocol described above. We used the pipeline described above to calculate the relative abundances of *Mycobacterium* (% of quality-filtered 16S rRNA gene sequences) on each of the swabs, after rarefying all samples to 8000 reads per sample. Twenty samples from North Carolina and all of the ‘negative control’ samples (both extraction blanks and no-template controls) were discarded due to insufficient sequencing depth (<1000 reads per sample), yielding a total of 64 swab samples included in downstream analyses. We used a PERMANOVA test (as implemented in the R package, vegan) to determine if storage duration had a significant effect on estimated mycobacterial abundances with sampling location (household) set as a fixed variable and unfrozen storage time as a random variable. We found no significant influence of storage time on the relative abundances of *Mycobacterium* in these samples (Pseudo-F = 1.67, P>0.20). Thus, we can conclude that potential differences in the duration of time samples spent unfrozen in transit are unlikely to have influenced our estimates of mycobacterial abundances on swabs collected as part of the broader sampling effort.

### Statistical analyses and predictive modeling

Of the 691 samples collected from across the U.S. and Europe, only 656 samples were included in downstream analyses. Thirty-five samples were excluded as corresponding information on household location (latitude/longitude), water source (municipal versus well water), frequency of showerhead usage, or showerhead material was not provided. Likewise, 37 of the 656 samples were excluded from the analyses of mycobacterial abundances (genus level) as insufficient 16S rRNA gene sequence data were available for those samples. For the water chemistry analyses, 45 of the 656 samples were excluded because no water chemistry data were provided by the citizen scientists.

We used permutational ANOVA (PERMANOVA) (67) to determine if there were significant differences in relative abundances of the genus *Mycobacterium* (16S rRNA gene sequence data) or individual mycobacterial lineages (hsp65 sequence data) across sample categories: household water source (municipal versus well), showerhead type, household location (U.S. versus Europe).

We next sought to identify whether there were spatial differences in the relative abundances of the genus *Mycobacterium* (16S rRNA gene sequence data) or individual mycobacterial lineages (hsp65 sequence data). As our samples were not collected randomly across the U.S. and Europe, but rather sampling intensity tended to track population density (with more samples collected from regions with larger cities), we first used hierarchical clustering via the hclust algorithm from the R “stats” package (https://cran.r-project.org) to group these samples into regional clusters based on their geographic proximity, excluding those clusters represented by fewer than 10 samples. Using these criteria, we ended up with 21 regional clusters of samples (shown in Figure 4 and Figures S8-S9) and the maximum distance between sampling locations within each cluster varied from 170-550 km and 200-510 km (east-west, and north-south distances, respectively). Eighty of the 656 samples did not fall into one of these 21 clusters, i.e. 80 samples were too spatially isolated to be assigned to any of the identified spatial clusters. For the 21 geographic clusters identified, we then used PERMANOVA analyses to determine if there were a significant difference in mycobacterial abundances across the regions. Spatial cluster was included as a fixed factor in these analyses. Similar analyses were conducted to determine if there were significant differences in measured water chemistry parameters across the 21 geographic regions.

We conducted semi-partial correlations (Spearman) using the ppcor R package (68) to evaluate the correlation between water chemistry data (total chlorine, free chlorine, pH, hardness, alkalinity, total iron, nitrite and nitrate) with the relative abundance of *Mycobacterium* (16S rRNA gene sequence data) and individual mycobacterial lineages (hsp65 sequence data). The mean annual temperature for each household location (from the Worldclim database worldclim.org) was also included in these analyses. Unlike standard correlations, semi-partial correlations allow us to identify the variance from a given response variable that is uniquely predictable from a given predictor, controlling for all other predictors simultaneously (69). We used a heatmap (heatmap.2 function in the R package gplots) to visualize our results.

Significant correlations (p<0.05) between the median mycobacterial abundance and prevalence estimates of pulmonary NTM disease generated from prior US population-level studies of Medicare beneficiaries (50) and persons with cystic fibrosis (70) were evaluated at the state-level for MAC and *M. abscessus.* These analyses were performed using SAS 9.4 (Carey, NC, USA).

## Acknowledgements

We want to thank Jessica Henley, Caihong Vanderburgh, Sarah McCoy, Robin Hacker-Cary, Julie Sheard, and Lea Shell for assistance with sample collection and processing. We also want to thank Ravleen Virdi for her help with the mycobacterial cultivation effort. Funding for this project was provided by the Innovative Research Program of the Cooperative Institute for Research in Environmental Sciences (N.F.), the High Plains Intermountain Center for Agricultural Health & Safety (N.F.), the U.S. Department of Defense (N.F., R.D.) and the Shoot for the Cure and Padosi Foundations (J.R.H). M.D-B. acknowledges support from the Marie Sklodowska-Curie Actions of the Horizon 2020 Framework Programme H2020-MSCA-IF-2016 under REA grant agreement n°70205

---

**Figure S1:**
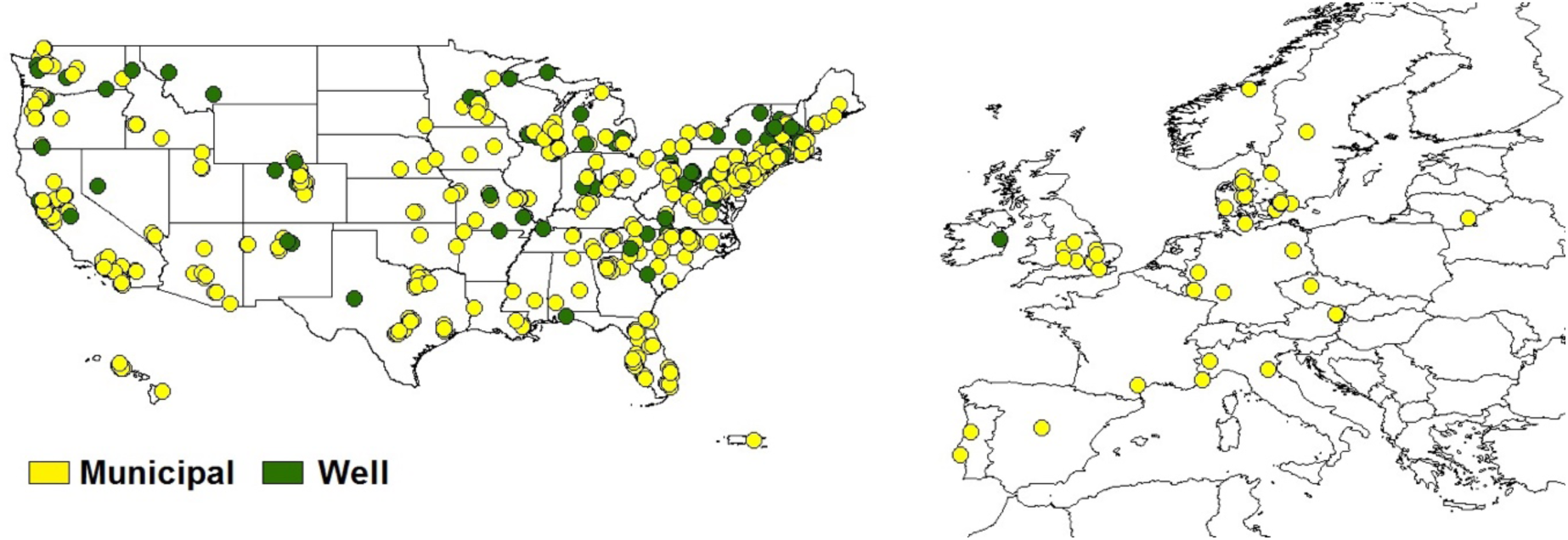
Map of the households from which showerhead biofilm samples were collected for this study (N = 656). The yellow points indicate households on municipal water (N = 520), while the green points indicate households on well water (N = 86).

**Figure S2:**
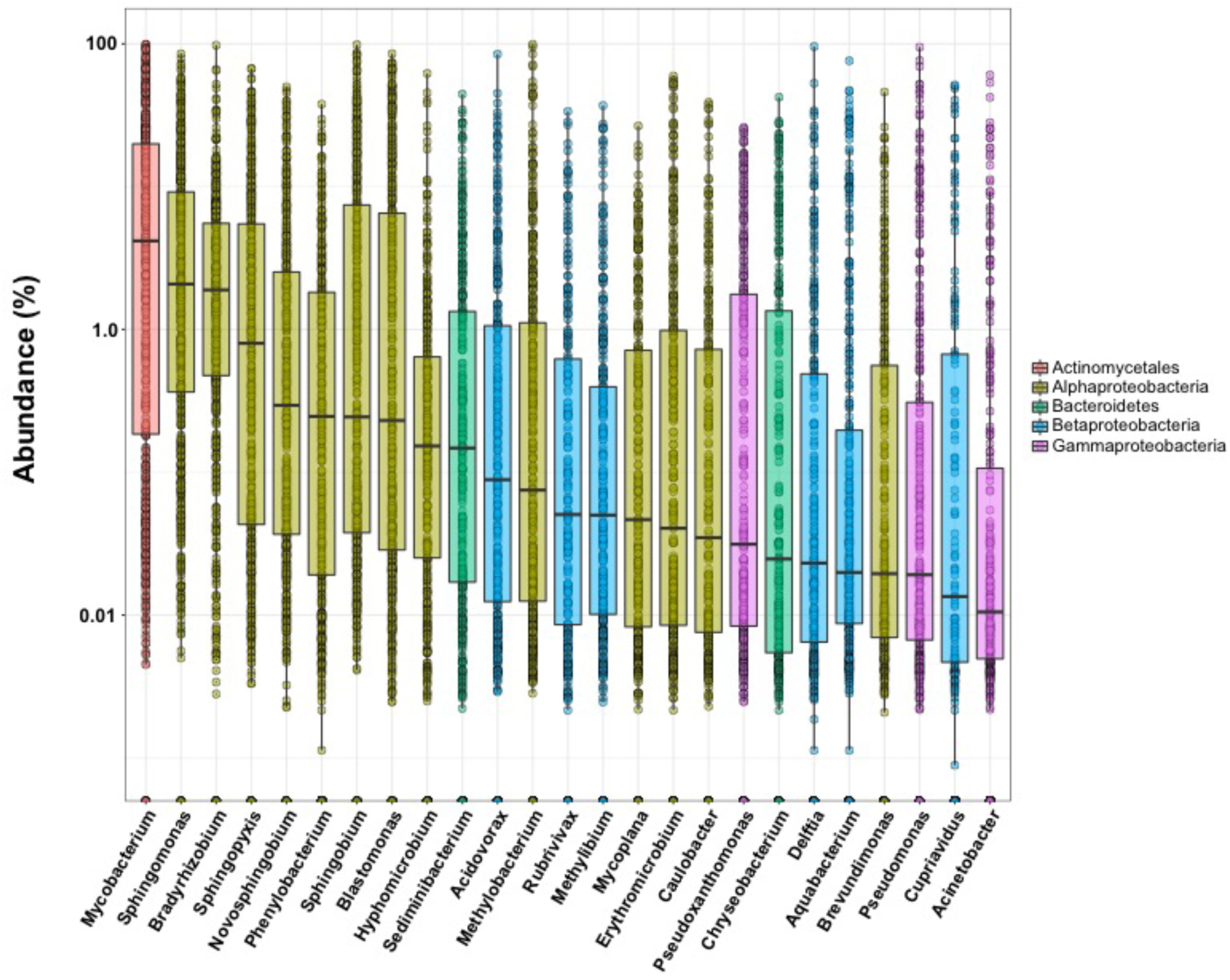
Box and whisker plots showing the dominant bacterial genera detected in 652 showerheads. Bars indicate mean abundances of each genus across all samples, the colors of each bar indicate the broader taxonomic affiliation of each genus. The y-axis shows the percentage abundances of each genus (log scale).

**Figure S3:**
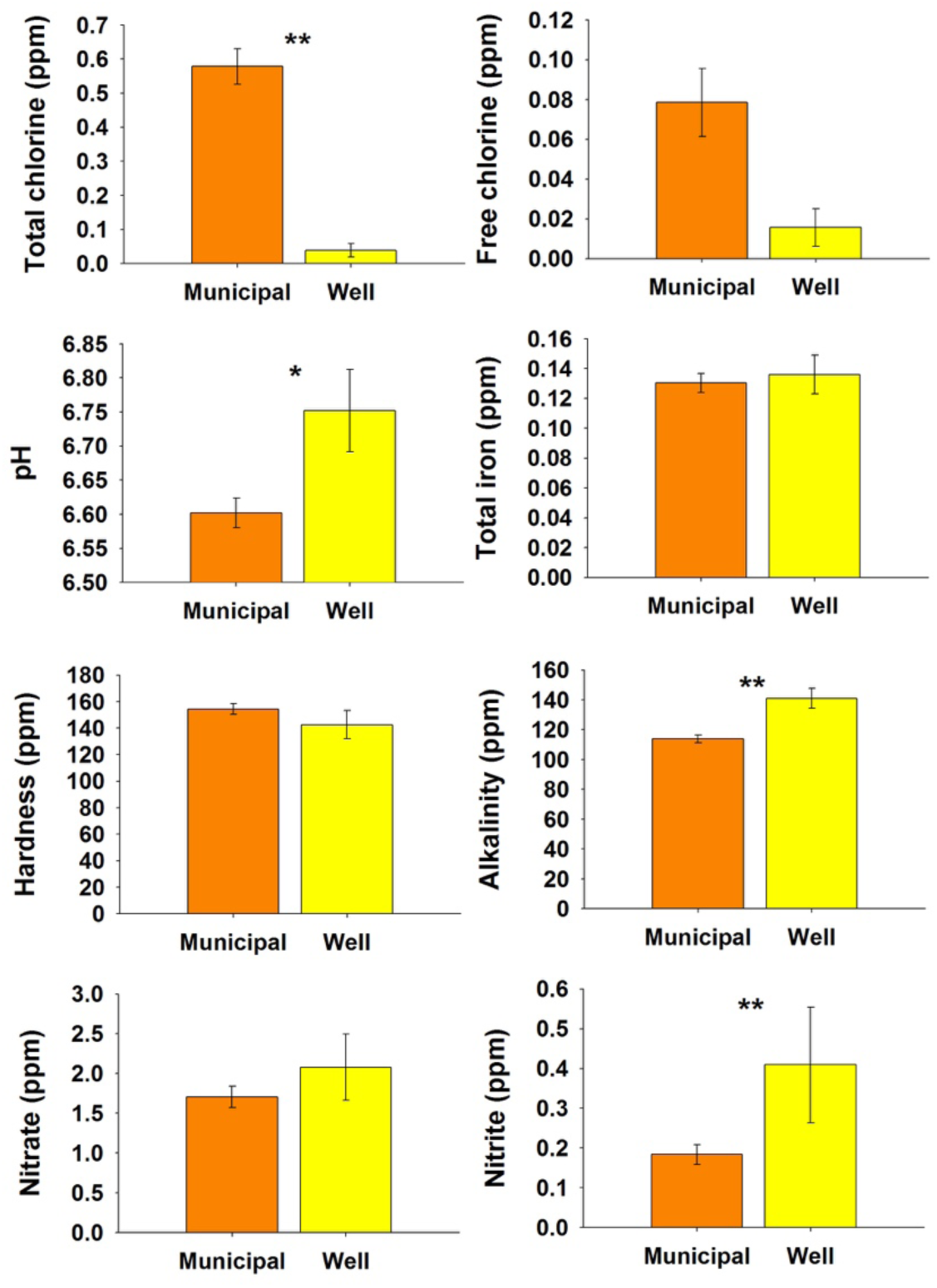
Differences in shower water chemistry across the U.S. households on either municipal or well water, as measured by the participants in this study.

**Figure S4:**
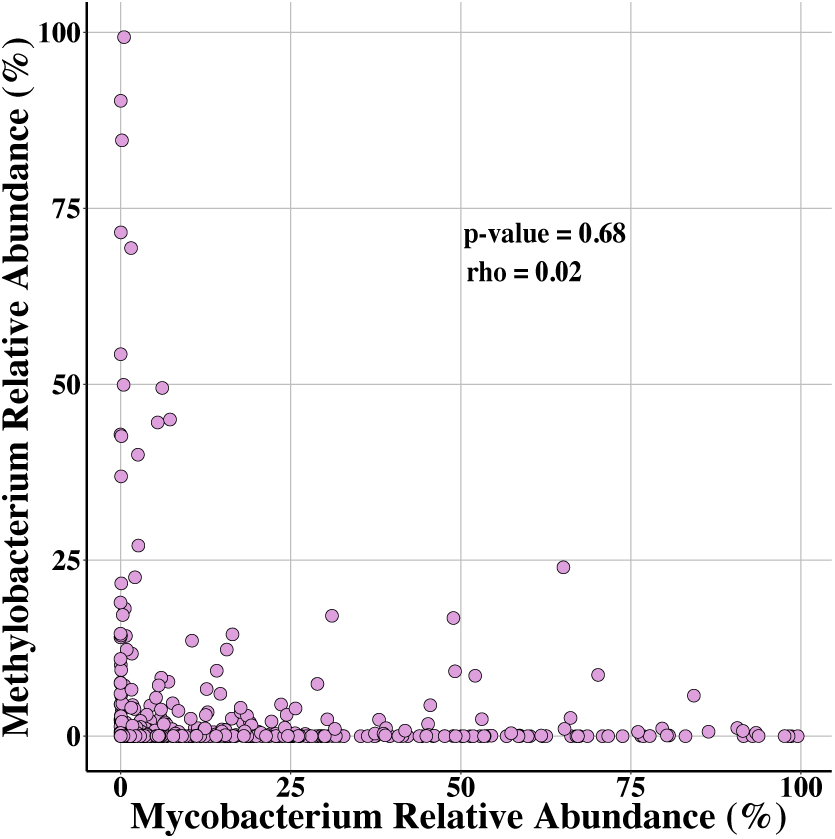
The relative abundance of *Methylobacterium* versus *Mycobacterium* across all samples for which we obtained 16S rRNA gene sequence data (N = 652).

**Figure S5:**
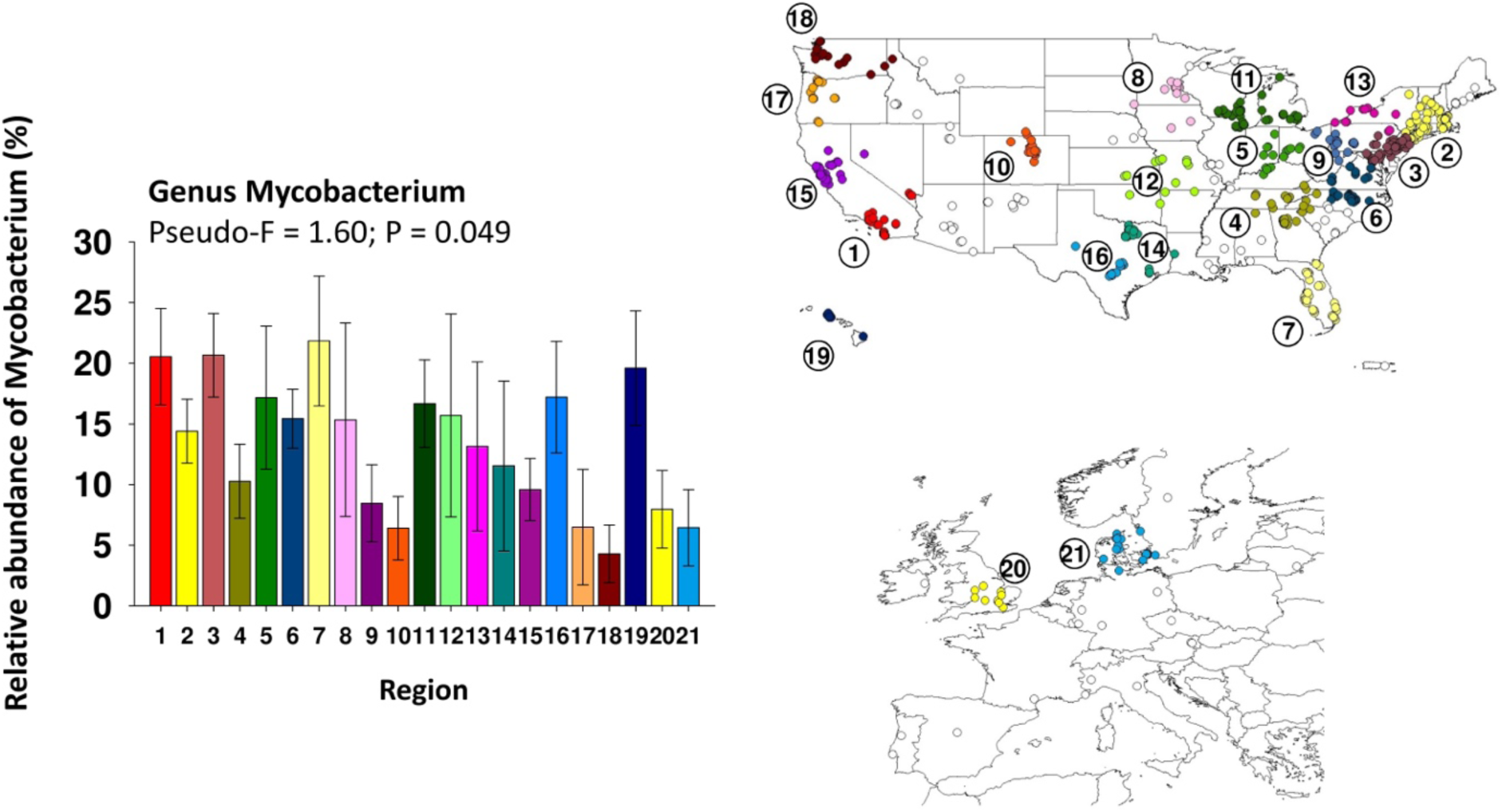
Relative abundance of the genus *Mycobacteria* across the 21 geographic clusters (regions) that had at least 10 samples per region. The different colored points on the map correspond to each of the 21 regions and the unfilled points indicate households that did not fall into one of these 21 geographic clusters.

**Figure S6:**
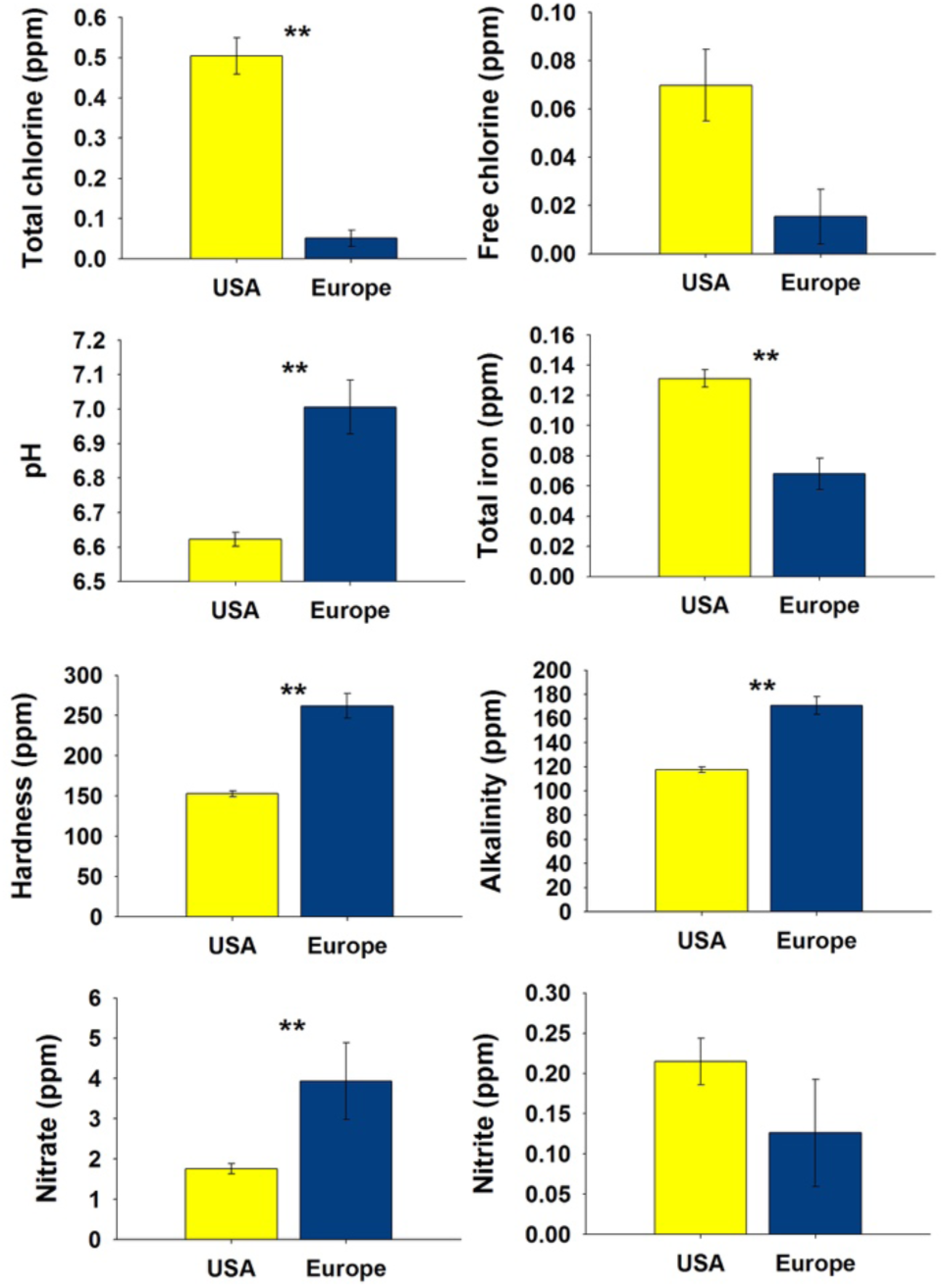
Differences in shower water chemistry across the U.S. versus European households. Only those households on municipal water supplies were included in these analyses (N=568 in total, 520 from the U.S., 48 from Europe).

**Figure S7:**
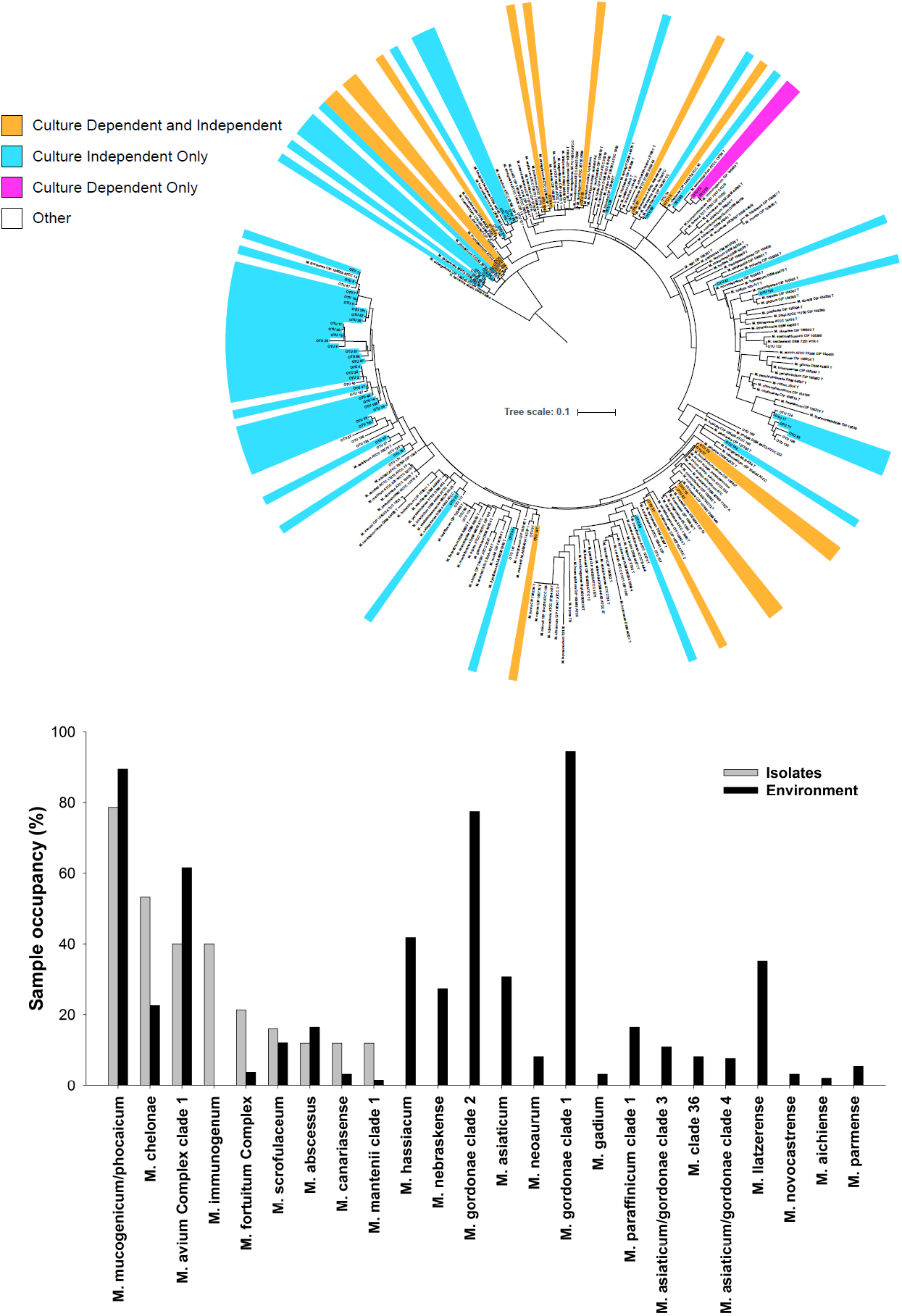
Phylogenetic tree indicating those hsp65 exact sequence variants (ESVs) detected in 186 samples for which we had conducted both cultivation-independent and cultivation-dependent assessments of mycobacterial diversity. Individual ESVs are colored by detection in both cultivation independent and dependent assays (orange), detection using the culture independent approach only (blue), and detection in culture only (pink). No color indicates either a database (reference) sequence or an ESV that was detected in another sample (not one of the 186 samples used for the culture independent versus dependent comparison). The bar plot below indicates the relative abundance of the dominant clades detected in the cultivation-independent analyses of DNA extracted directly from the environmental samples (showerhead biofilms −black bars) versus analyses of the DNA extracted from the cultivated mycobacterial isolates (grey bars).

**Figure S8:**
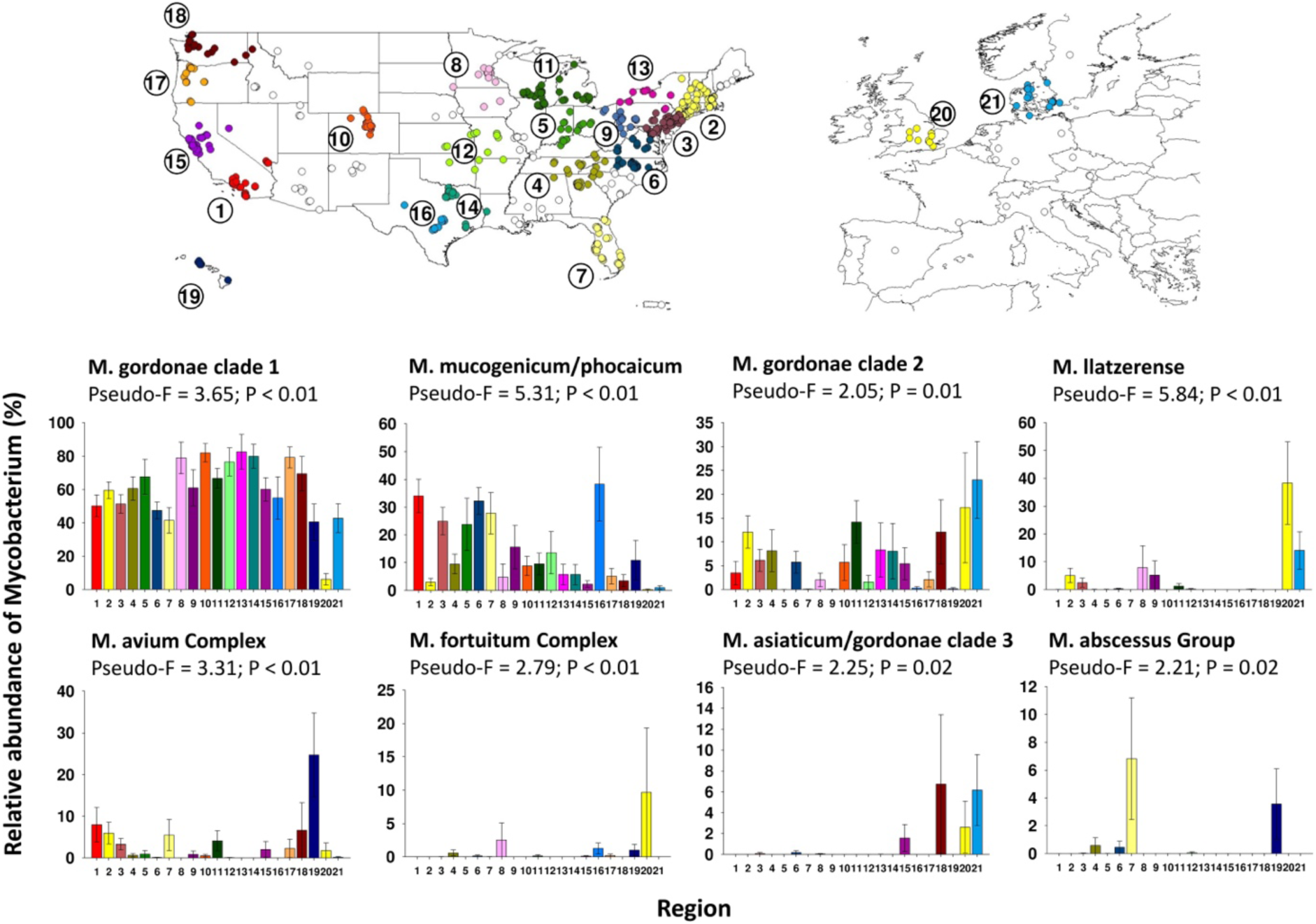
Relative abundance of mycobacterial clades across the 21 geographic clusters (regions) that had at least 10 samples per region. The different colored points on the map correspond to each of the 21 regions and the unfilled points indicate households that did not fall into one of these 21 geographic clusters. Only those clades that exhibited significant differences in relative abundance across regions (P<0.05) are shown here. For information on the geographic distributions of the remaining abundant clades, see Figure S9.

**Figure S9:**
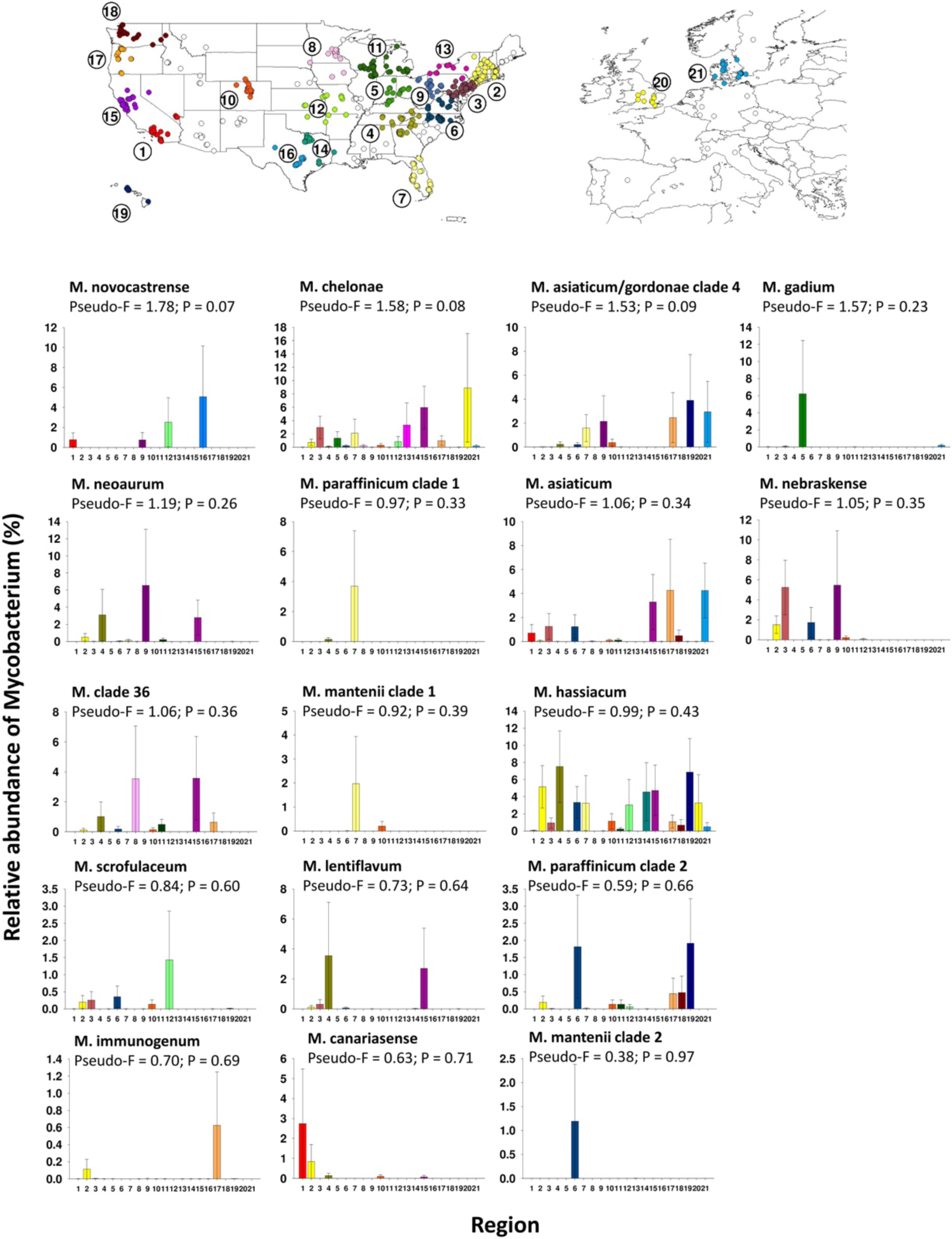
Relative abundance of the remaining mycobacterial clades (those not shown in Figure S8) across the 21 geographic clusters (regions) that had at least 10 samples per region. The different colored points on the map correspond to each of the 21 regions and the unfilled points indicate households that did not fall into one of these 21 geographic clusters. For the 17 clades shown here, none exhibited statistically significant (P<0.05) variation in abundances across the 21 geographic clusters.

**Figure S10:**
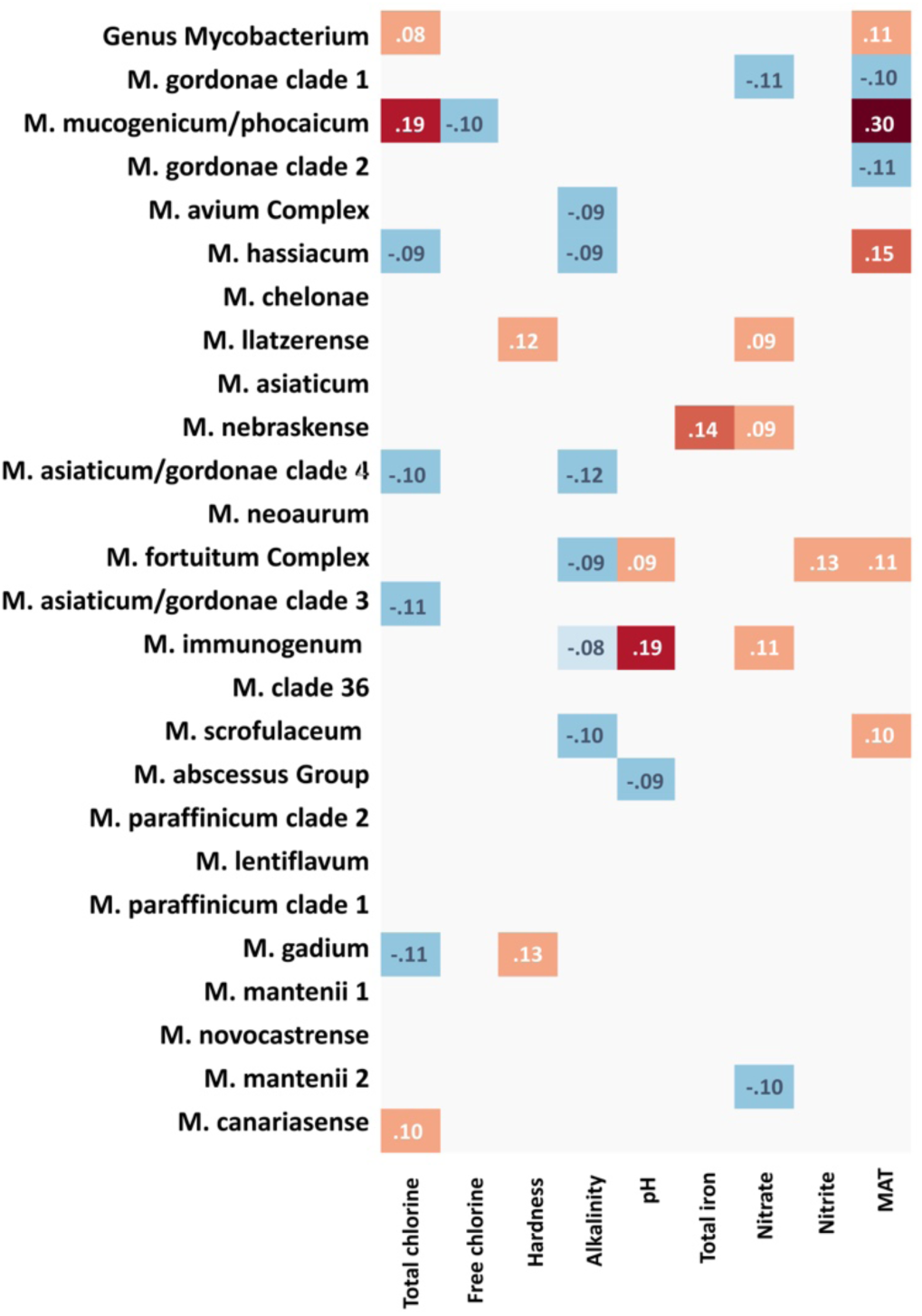
Semi-partial correlations (Spearman) indicating the correlations between water chemistry parameters and mean annual temperature (‘MAT’) with the relative abundance of mycobacteria (16S rRNA gene sequence data) and individual mycobacterial clades (hsp65 sequence data).

**Figure S11:**
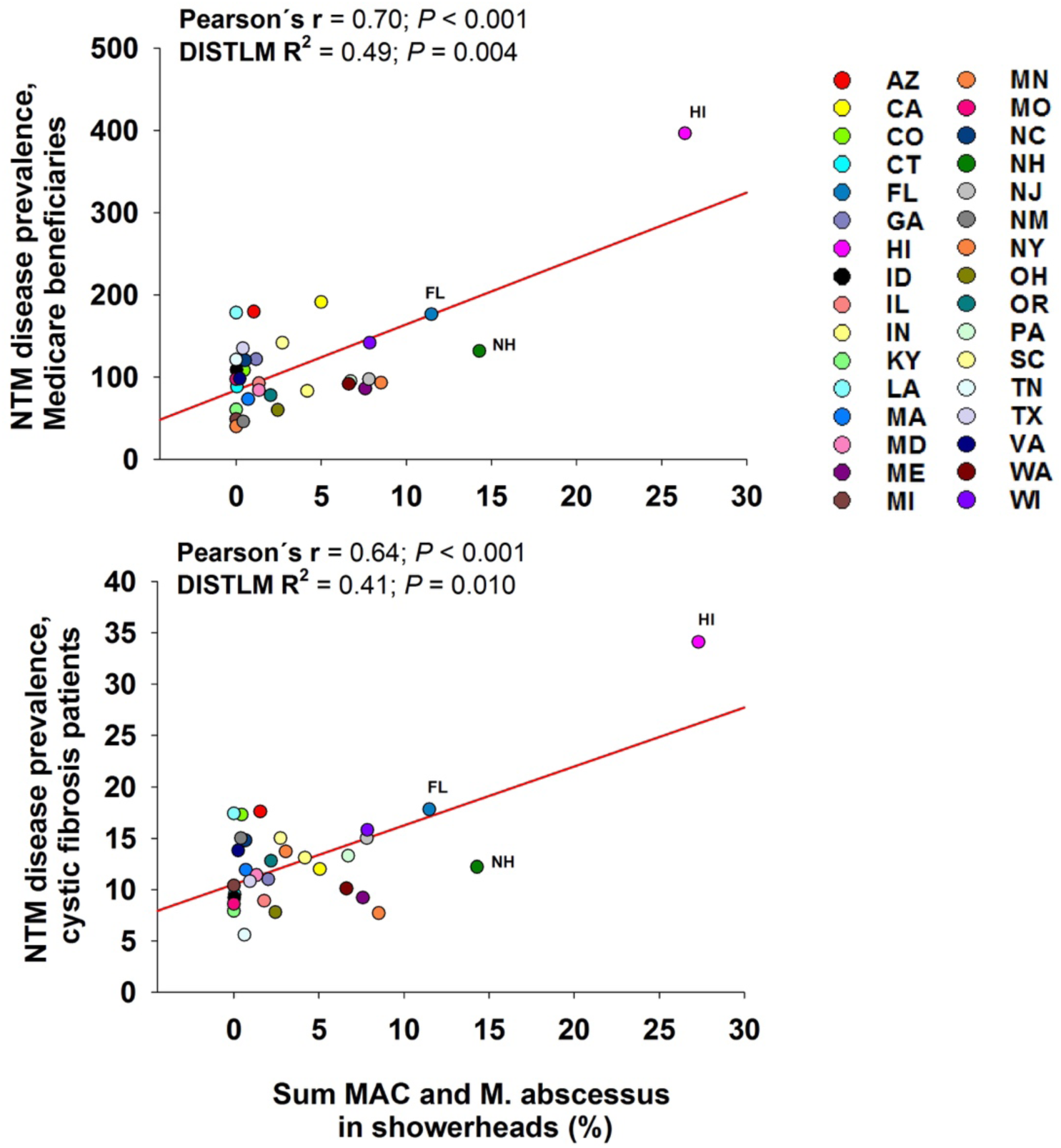
Relationships between the summed relative abundances of three potentially pathogenic mycobacterial clades that were frequently detected in showerheads (*M. abscessus, M. fortuitum*, and MAC) and the reported prevalence of NTM disease across Medicare beneficiaries and cystic fibrosis patients (50, 70). The strength and significance of the correlations were tested using both Pearson correlations and distance-based linear models (DISTLM, output shown). Each point indicates a different U.S. state and data were aggregated to the state level (using median abundances) as the disease prevalence data were only available at the state level of resolution. Only states with >10 showerhead samples were included in these analyses.

